# Suppression of FUS aggregation by proteasome inhibitors

**DOI:** 10.1101/2024.09.17.613412

**Authors:** Amal Younis, Kanar Yassen, Kinneret Rozales, Tahani Kadah, Naseeb Saida, Anatoly Meller, Ophir Lorian, Nadeen Akaree, Ronit Heinrich, Flonia Levy-Adam, Shai Berlin, Reut Shalgi

## Abstract

ALS (Amyotrophic Lateral Sclerosis) is characterized by aggregation of RNA-binding proteins, including FUS. The proteasome generally targets misfolded proteins for degradation, thereby limiting protein aggregation. Strikingly, we discovered that proteasome inhibitor treatment led to a significant suppression of ALS-associated mutant FUS aggregation in cells and primary neurons, in sharp contrast to its aggregation-promoting effects on other neurodegeneration-associated proteins. Our data revealed that proteasome inhibitors stimulated a unique transcriptional program, that selectively suppressed FUS aggregation. A key mediator of this program was the HSP70 co-chaperone BAG3, which markedly reduced FUS aggregation through selective autophagy. In parallel, the polyamine N1-acetyltransferase SAT1 was massively induced as part of this program, accompanied by remodeling of polyamine metabolism, and its metabolic activity was required for FUS aggregation suppression. Together, these findings reveal an unexpected protective response to proteasome inhibition that selectively suppresses FUS aggregation, which could be harnessed for future therapeutic intervention in FUS-ALS.

## Introduction

Pathological aggregation is a hallmark of neurodegenerative diseases^1, 2^. ALS (Amyotrophic lateral sclerosis), a devastating neurodegenerative disease with no effective treatment or cure, is characterized by protein aggregates in motor neurons. RNA-binding proteins (RBPs) have been shown to aggregate in ALS, with TDP-43 and FUS representing the most well-characterized examples^3, 4, 5^. In addition, mutations in numerous RBPs have been identified as causative of familial ALS^6^.

The protein quality control (PQC) network, composed of molecular chaperones and protein degradation pathways, including the UPS (Ubiquitin Proteasome System) and autophagy, maintains protein homeostasis under fluctuating physiological and environmental conditions^7, 8, 9, 10^. While the UPS sustains protein turnover under homeostasis, it further acts to specifically degrade damaged proteins upon stress, in cooperation with the chaperone network^10, 11, 12^. Nevertheless, in neurodegenerative diseases, the PQC network fails to prevent aggregation^8^. The role of the PQC network in regulating ALS-associated protein aggregation has been demonstrated previously^13, 14, 15, 16, 17, 18^, with accumulating evidence suggesting that distinct ALS drivers require subtype-specific PQC mechanisms to effectively combat aggregation^17, 18^.

Several proteostasis-related compounds have been investigated as potential therapies for ALS in recent years. Although some have shown considerable promise in animal models and small-scale subtype specific studies^19, 20^, they largely failed phase 2/3 clinical trials^21, 22^. These findings further suggest that, while proteostasis modulators retain considerable therapeutic promise, identifying the proteostasis requirements of distinct ALS drivers will be essential for selecting subtype-specific, effective therapeutic interventions.

Here we discovered an unexpected proteostasis modulator of FUS aggregation, namely, proteasome inhibitors. Importantly, while proteasome inhibitors are known to promote the accumulation of misfolded proteins due to impaired degradation, thereby aggravating pathological aggregation^10, 23, 24^, we found that proteasome inhibitors of various types significantly reduced FUS aggregation in cell line models and in primary neurons. This effect was primarily mediated through a unique transcriptional program, that was beneficial for FUS aggregation prevention. We focus on two prominent mediators of this program: the HSP70 co-chaperone BAG3, and the polyamine N1-acetyltransferase SAT1, further revealing a novel effect of proteasome inhibitors on polyamine metabolism. While proteasome inhibitors are FDA-approved drugs, our data positions BBB-permeable proteasome inhibitors, such as Marizomib^25^, could be therapeutically beneficial in FUS-ALS.

## Results

### Proteasome inhibitor treatment significantly reduces FUS aggregation

Previously, we identified specific chaperones that can potently reduce the aggregation of ALS-associated mutant FUS^17^. We next sought to further investigate the role of the protein quality control (PQC) network in modulating protein aggregation, and specifically, the potential role of the proteasome. We first employed MG-132, a direct proteasome inhibitor^26^, which leads to the accumulation of misfolded proteins due to impaired degradation, and measured its effects on aggregation. Based on previous studies, we expected proteasome inhibition to promote protein aggregation, as reported in the literature for several neurodegeneration-related proteins, such as HTT-polyQ^23^, TDP-43^24^ and others^10, 12, 27^.

We treated HTT-polyQ cell models with MG-132 for 18 hours, and measured the change in the percentage of aggregate-containing cells using the FACS-based PulSA assay^28^, which is able to distinguish aggregate-containing from non-aggregate cells based on their fluorescent signal properties (Fig. S1A). The analysis produces an aggregation modulation score, defined as the log2 fold change in the percentage of aggregate positive (AGG+) cells in treated cells compared to the control^17^ (Fig. S1A,B see Methods). Our results confirmed the anticipated exacerbating effect of proteasome inhibitors on HTT-polyQ aggregation, consistent with previous literature^23^ (Fig. 1A, S1B,C). Similarly, in cells expressing an aggregation prone variant of TDP-43 (TDP-43-△NLS-2KQ^29^), MG-132 treatment significantly enhanced TDP-43 aggregation (Fig. 1B, S1B,D). Surprisingly, however, when subjected to identical conditions of MG-132 treatment, cells expressing FUS with an ALS-related mutation R521H (mutFUS) exhibited an unexpected reduction in aggregation by more than 30% compared to control (Fig. 1C,D, S1B,E,F). Under these conditions, there was no significant impact on cell viability (Fig. S1G). Importantly, the effects on mutFUS aggregation, as well as on TDP-43-△NLS-2KQ and HTT-polyQ aggregation, were independently confirmed by fluorescence microscopy imaging and quantitative image analysis (Fig. S1C-E,H).

**Figure 1:**
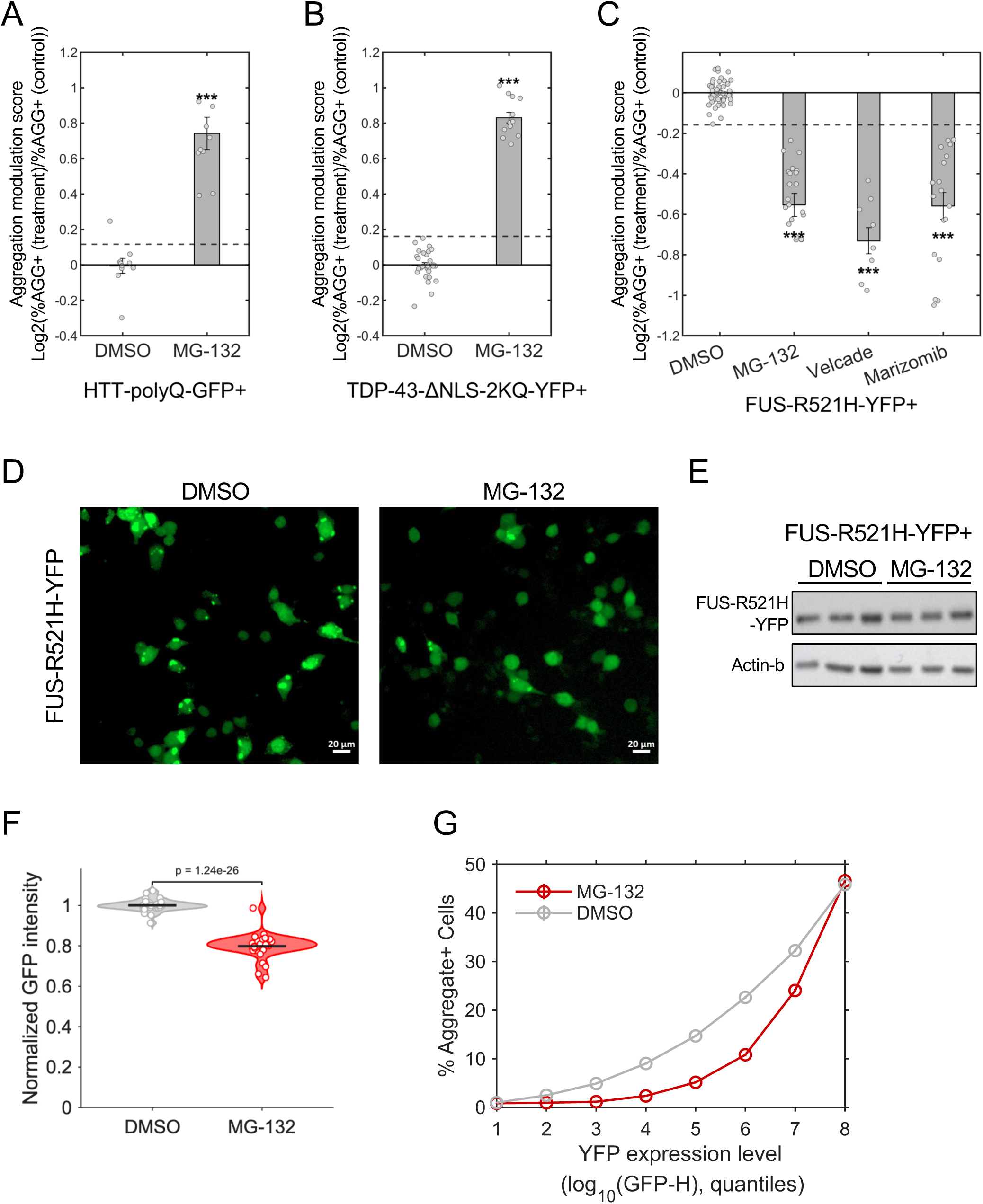
Proteasome inhibitor treatment significantly reduced FUS aggregation. (A-B) Aggregation modulation scores for cells expressing HTT-polyQ (A), or TDP-43-ΔNLS-2KQ (B) demonstrating an exacerbating effect of MG-132 treatment on aggregation. Data presented as mean values ± SEM of N = 3 independent biological experiments, with a total of n = 10/13 replicate samples for HTT/TDP-43, respectively. (C) FUS-R521H-YFP aggregation modulation scores for cells expressing mutFUS treated with various proteasome inhibitors (see Methods), including MG-132, Velcade (Bortezomib), and Marizomib, all of which exhibit unexpected significant aggregation suppression. Data presented as mean values ± SEM of N = 8/3/6 independent biological experiments with a total of n = 22/9/18 replicates for MG-132/Velcade/Marizomib respectively. (A-C) Dashed lines represent 95% CIs. (HTT CI = +/-0.1168, TDP-43 CI = +/-0.1612, FUS CI = +/- 0.1577). ***p < 0.003, *p < 0.05, empirical p-value (see Methods). (D) FUS-R521H-YFP fluorescence microscopy images, with control (left) and MG-132-treated (right, 10μM for 18h) cells. Images are representative N=4 independent experiments (see S1Gnew for quantification by image analysis). Scale bars - 20 μm. (E) HEK293T cells expression FUS-R521H-YFP were treated with either 10 µM MG-132 for 18 h or DMSO as a control. FUS-R521H-YFP (∼95 kDa) protein levels were assessed by immunoblotting using an anti-GFP antibody, with actin (∼42 kDa) serving as a loading control. Densitometric quantification revealed no significant difference in FUS-R521H-YFP protein levels between MG-132 and DMSO treated cells (Fig. S1I). (F) Violin plots display the distributions of GFP fluorescence in YFP+ HEK293T cells expressing mutant FUS-R521H-YFP following DMSO or MG132 treatment, as measured by flow cytometry. Horizontal lines denote mean values. (G) FUS-R521H-YFP aggregation suppression by MG-132 is robust across FUS-R521H-YFP expression levels. HEK293T cells expressing FUS-R521H-YFP were treated with MG-132 or DMSO as control. Raw flow cytometry data were binned according to FUS-YFP level and the proportion of cells with aggregates (AGG⁺ population) from the total number of cells in each bin was plotted on the y-axis (see Methods). Data presented as median ± SEM, n=∼361,000/372,000 for DMSO/MG-132 treated cells respectively.

Although western blot analysis revealed no significant change in mutFUS protein levels following MG-132 treatment (Fig. 1E, S1I), flow cytometry detected a modest, ∼20% reduction in FUS-R521H-YFP fluorescence intensity (Fig. 1F). Still, suppression of mutFUS aggregation was consistently observed across the entire range of mutFUS expression levels, demonstrating that the anti-aggregation effect of MG-132 is not merely a consequence of reduced protein levels (Fig. 1G).

To determine whether this unexpected aggregation suppression was restricted to MG-132, we evaluated the impact of additional proteasome inhibitors. Treatment with Velcade (Bortezomib), an FDA-approved therapeutic agent for multiple myeloma, or Marizomib, a proteasome inhibitor that was shown to cross the BBB (Blood Brain Barrier)^25^, similarly resulted in a significant reduction in mutFUS aggregation (Fig. 1C, S1F,J-L). Moreover, the suppression of FUS aggregation by different proteasome inhibitors was recapitulated for additional ALS-associated FUS mutations (Fig. S1M,N).

### Proteasome inhibitor-induced suppression of FUS aggregation is transcriptionally mediated

We next investigated the effects of an additional PQC modulator, Withaferin A, a steroidal lactone from the medicinal plant *Acnistus arborescens*, which was previously identified as an activator of HSF1^30^, the Heat Shock response transcription Factor^31^. Cells were exposed to Withaferin A for 18h, and subsequent PulSA analysis showed that Withaferin A significantly alleviated mutFUS aggregation (Fig. 2A). This result prompted us to examine the potential involvement of HSF1, and the proteotoxic stress response, downstream of proteasome inhibitors as a mediator of mutFUS aggregation repression. Indeed, proteasome inhibitor treatment is known to trigger a transcriptional program similar to the proteotoxic stress response mediated by HSF1^32^. To determine whether the mechanism underlying proteasome inhibitors suppression of mutFUS aggregation is HSF1 dependent, we utilized small interfering RNA (siRNA) targeting HSF1. siHSF1 reduced HSF1 endogenous levels by at least 70% (Fig. S2A). Cells co-transfected with FUS-R521H-YFP and siControl exhibited a significant mutFUS aggregation suppression upon MG-132 treatment (Fig. 2B). In contrast, treatment with MG-132 in siHSF1 expressing cells failed to alleviate mutFUS aggregation (Fig. 2B). Notably, induction of the heat shock response by heat treatment did not reduce mutFUS aggregation, as we reported previously^17^ (Fig. S2B), yet silencing of HSF1 impaired the ability of MG-132 to suppress mutFUS aggregation (Fig. 2B). We therefore hypothesized that proteasome inhibitors induce a unique transcriptional program, distinct from that elicited by heat shock, which is specifically beneficial against FUS aggregation.

**Figure 2:**
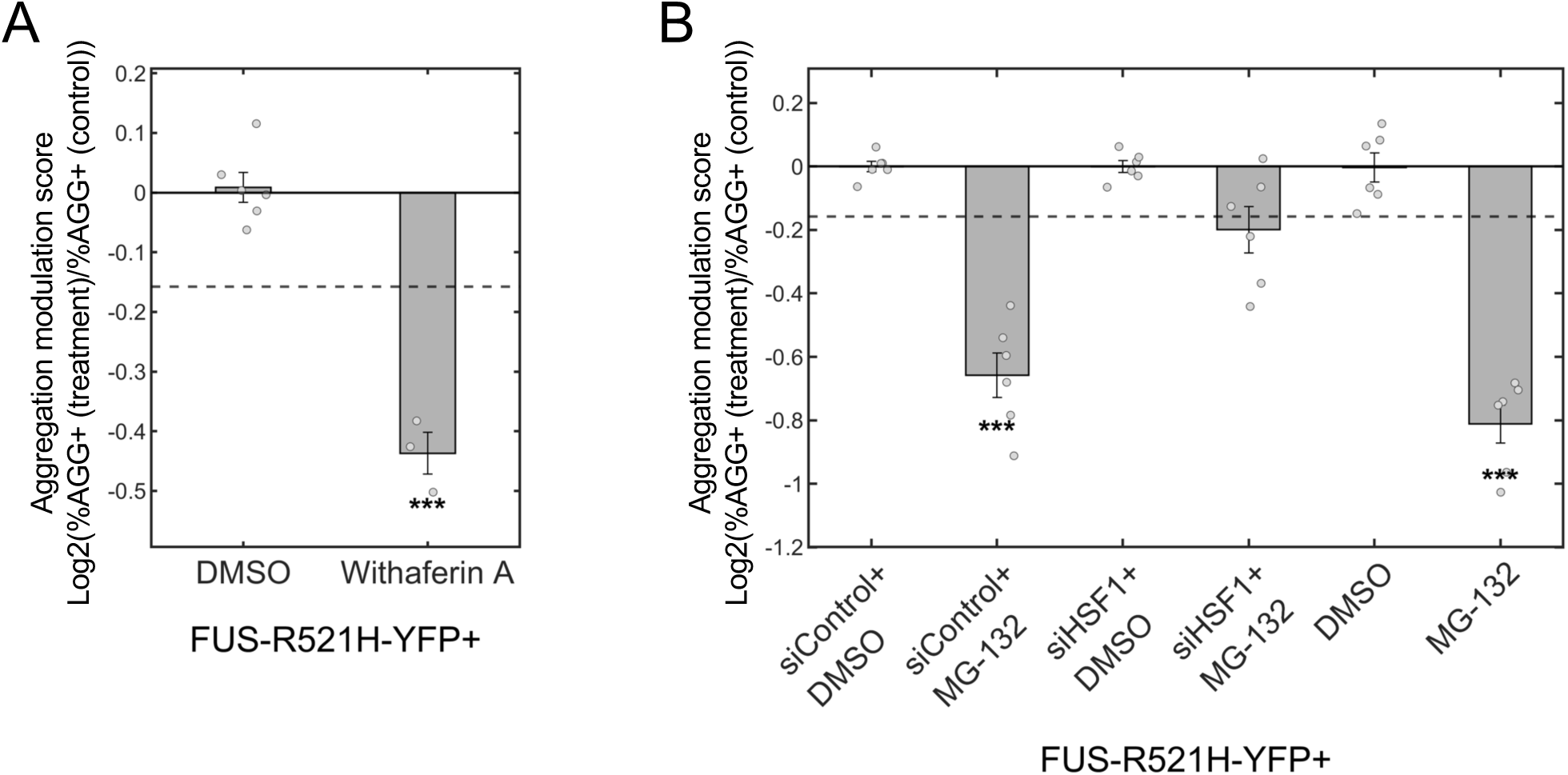
Proteasome inhibitor suppression of FUS aggregation is HSF1 dependent. (A) PulSA analysis of cells expressing FUS-R521H-YFP treated with 6 µM Withaferin A (WA) for 18 h demonstrates suppression of FUS aggregation compared to DMSO treated controls. Data presented as mean ± SEM of N = 3 independent biological experiments, a total of n = 6/3 replicate samples for DMSO/WA, respectively. (B) FUS-R521H-YFP Aggregation modulation scores in cells expressing mutFUS that underwent HSF1 knockdown (siHSF1) vs. siControl cells in addition to cells expressing FUS-R521H-YFP without siRNA. Cells were treated with MG-132 10uM for 18h and subjected to PulSA analysis. Results show that the suppressive effects of MG-132 on FUS aggregation are dependent on HSF1. Data presented as mean ± SEM from N = 3 independent biological experiments, with a total of n = 6 replicate per condition. (A,B) Dashed lines represent 95% CIs ***p < 0.003, empirical p-value (see Methods).

### The chaperone BAG3 is a key mediator of the suppressive effects of proteasome inhibitors on FUS aggregation

We next sought to identify genes involved in the proteasome inhibitors-mediated transcriptional program. We therefore performed an RNA-seq experiment, treating cells with the proteasome inhibitor MG-132, or with heat shock, for different durations. We reasoned that genes mediating the suppressive effect of proteasome inhibitors on FUS aggregation would be induced to a greater extent by MG-132 than by heat shock. We therefore looked at the change (log2 fold change, LFC) in gene expression in cells treated with MG-132 (normalized to their respective control samples), compared to the change in cells subjected to heat shock (normalized to their respective controls). Analysis of PQC-related pathways showed that chaperones and ubiquitin ligases were significantly induced by both treatments (Fig. 3A). Interestingly, genes related to core autophagy, as well as autophagy regulators, were selectively induced in response to MG-132 but not in response to heat shock (Fig. 3A). A closer examination of these genes showed that, while the changes in expression of all these genes combined were largely correlated between proteasome inhibitors and heat shock treatments (Fig. 3B, pearson r=0.61, p=1e-43), several mRNAs were induced to a higher extent in the presence of MG-132 compared to heat shock (Fig. 3B). Those included a few transcription factors, and several canonical HSP70 genes, which play a generalized role in proteostasis (Fig. S2C). Other more specific factors included the chaperones DNAJB1, CRYAB, and BAG3. Functional analysis using PulSA showed that, while DNAJB1 and CRYAB did not affect FUS aggregation (Fig. S2D), the HSP70 co-chaperone BAG3, which is also known as an autophagy regulator^33^, was found to significantly reduce FUS aggregation (Fig. 3C, S3A, S1F)^17^.

**Figure 3:**
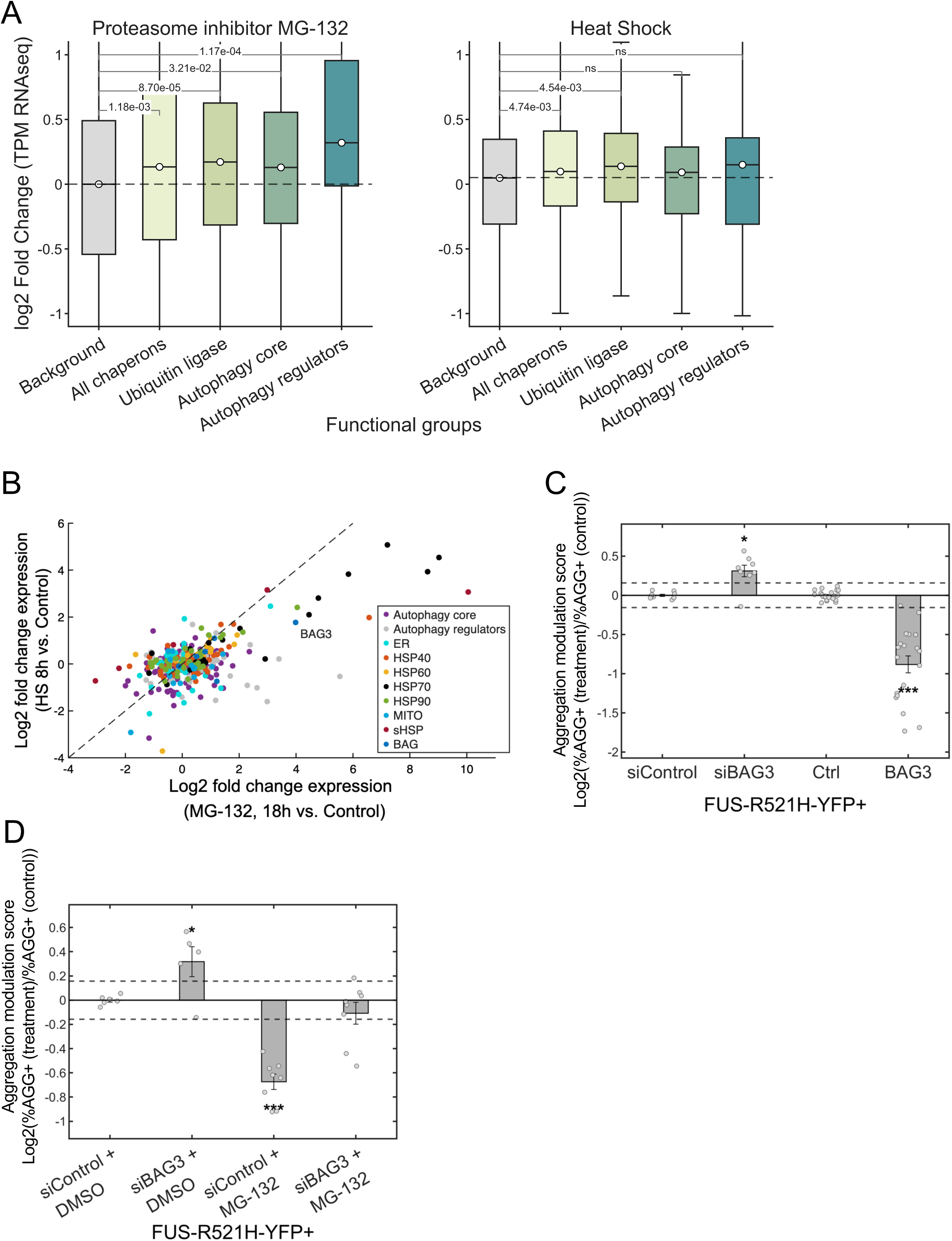
BAG3 is a key mediator of the repressive effects of proteasome inhibitor on FUS aggregation. (A) Gene expression changes of protein quality control (PQC) related gene groups following MG-132 treatment (10uM, 18h, left) or heat shock (8h, 42°C, right). Box plots show the distribution of log2 fold changes in gene expression relative to respective controls. Chaperones and ubiquitin ligases were significantly induced by both treatments, whereas genes involved in core autophagy or autophagy regulation were selectively induced following MG-132 treatment but not heat shock. FDR-adjusted p-values are indicated above the plots. (B) Log2-fold changes (LFC) of gene expression for cells treated with MG-132 (10uM for 18h) compared to control (x-axis) plotted against the LFC of genes in cells subjected to heat shock (42°C for 8h) compared to their respective controls (y-axis). Diagonal - dashed line. Functional group genes are color coded. (C) mutFUS Aggregation modulation scores in cells expressing FUS-R521H-YFP that underwent BAG3 knockdown (siBAG3) vs. siControl, alongside cells co-expressing FUS-R521H-YFP with either DsRed (Ctrl) or BAG3. BAG3 markedly reduces mutFUS aggregation, while BAG3 knockdown significantly increases aggregation. Data presented as mean ± SEM of at least N=3 biologically independent experiments, comprising n = 9, 8, 22, and 19 total replicates for siControl, siBAG3, DsRed, and BAG3, respectively. (D) mutFUS aggregation modulation scores in cells expressing FUS-R521H-YFP that underwent BAG3 knockdown or siControl, and were treated with either MG-132 or DMSO, showing that aggregation suppression by MG-132 is dependent on BAG3. Data presented as mean ± SEM of N = 3 biologically independent experiments, with a total of n=6/5/8/8 replicates for siControl+DMSO/siBAG3+DMSO/siControl+MG-132/siBAG3+MG-132, respectively. (B-D) Dashed lines represent 95% CIs ***p < 0.003, *p < 0.05, empirical p-value (see Methods).

While BAG3 significantly reduced mutFUS aggregation, it did not colocalize with mutFUS aggregates (Fig. S3B). To further investigate the inhibitory role of BAG3 on mutFUS aggregation, we used siRNAs to silence BAG3 expression, which led to a 97% reduction in its endogenous levels (Fig. S3C). We then used FACS to assess its impact on aggregation. The results demonstrated a significant exacerbation of aggregation following BAG3 knockdown (Fig. 3C), underscoring the critical role of BAG3 in mutFUS aggregation regulation. This effect was specific to mutFUS, as siBAG3 reduced TDP-43-△NLS-2KQ aggregation, while overexpression of BAG3 promoted TDP-43-△NLS-2KQ aggregation, in agreement with a recent report^34^ (Fig. S3D).

Next, we treated mutFUS expressing cells, transfected with either siBAG3 or siControl, with MG-132 or DMSO as a control (Fig. 3D). Importantly, the suppression of mutFUS aggregation by MG-132 was completely lost in the absence of BAG3 (Fig. 3D). These findings identify BAG3 as an essential mediator of the MG-132-induced transcriptional program that suppresses mutFUS aggregation.

### BAG3 inhibits FUS aggregation through selective autophagy

BAG3, a nucleotide exchange factor (NEF), contains a conserved C-terminal BAG domain, which binds the ATPase domain of HSP70^35^ and facilitates the release of HSP70 clients to complete a chaperoning cycle^35, 36^. To investigate whether the inhibitory effect of BAG3 on mutFUS aggregation depends on HSP70, we generated a BAG3 isoform lacking the C-terminal BAG domain (BAG3-ΔC, Fig. 4A). We then tested the interaction of BAG3 or of BAG3-ΔC with HSP70 via co-immunoprecipitation (co-IP), and showed that while the full-length BAG3 interacted with HSP70, the C-terminal truncation completely abrogated this interaction (Fig. 4B, S4A,B). When employing the PulSA assay on cells co-expressing FUS-R521H-YFP and BAG3-ΔC, we found that this mutant caused a slight but significant enhancement of FUS aggregation (Fig. 4C).

**Figure 4:**
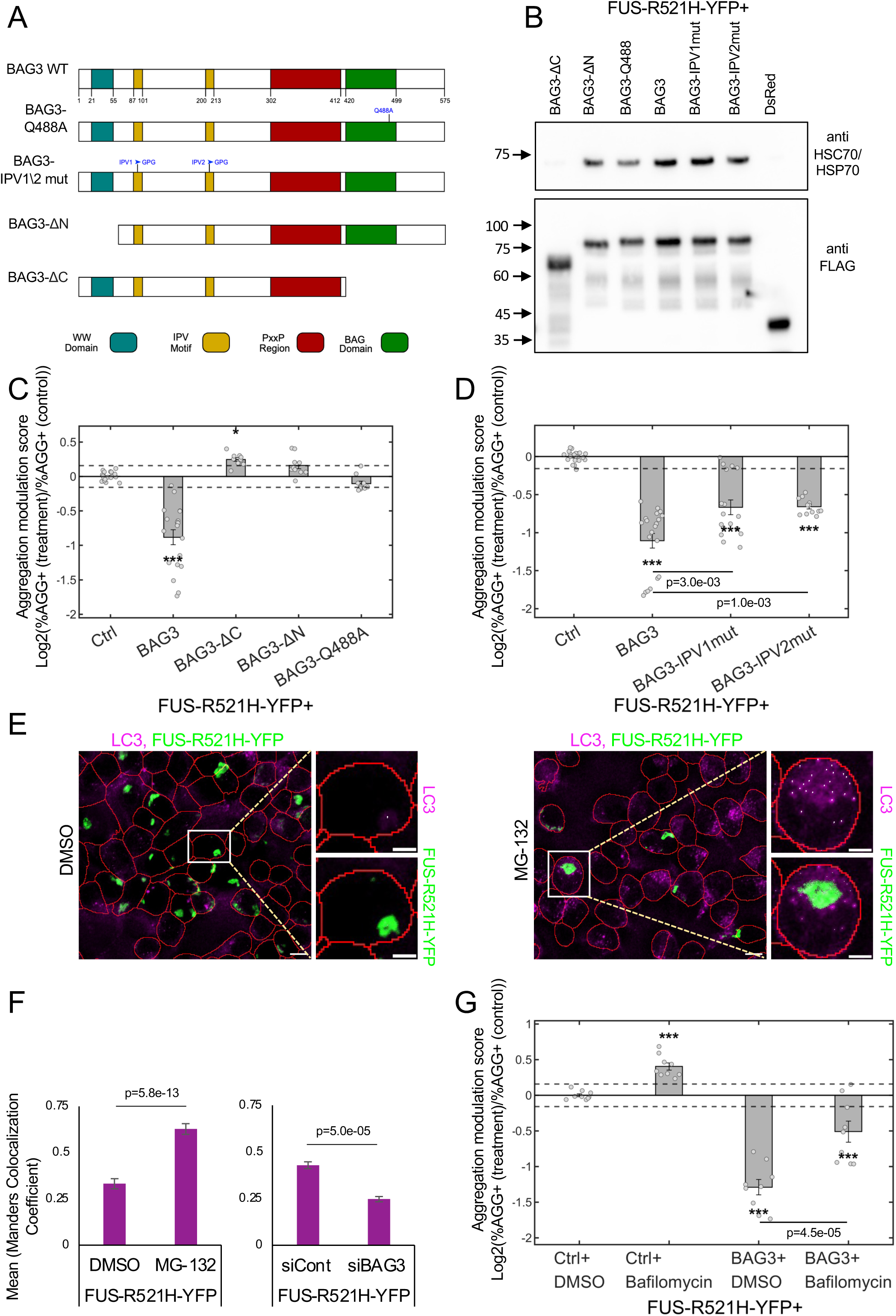
BAG3 inhibits FUS aggregation through selective autophagy. (A) BAG3 isoforms used in this study. BAG3 consists of 575 amino acids with a conserved C-terminal BAG domain facilitating HSP70 binding. BAG3-ΔC: C-terminally truncated BAG3 isoform. Q488A: point mutation in the BAG domain. BAG3-ΔN: N-terminal truncated BAG3 isoform lacking the WW domain,. IPV1/2mut: BAG3 mutated for the two conserved IPV (Isoleucine-Proline-Valine) motifs, known to be important for binding of sHSPs. (B) Co-IP for different BAG3 isoforms (FLAG-tagged) with the endogenous HSC70/HSP70 showed no interaction with BAG3-ΔC or DsRed and reduced interaction with Q488A. (See Fig. S4A,B, for quantification and inputs). The expected molecular weight of full-length BAG3 in the 3xFLAG-V5 plasmid is 69kDa, BAG3-ΔC is ∼52, and BAG3-ΔN is ∼62 kDa. (C) FUS-R521H-YFP Aggregation modulation scores for BAG3 isoforms, showing that BAG3 mutants lose their repressive effects on FUS aggregation. Data presented as mean ± SEM of N = 5 biologically independent experiments, with a total of n = 10 replicates for each isoform and n = 19 replicates for full-length BAG3. (D) FUS-R521H-YFP aggregation modulation scores for either BAG3-WT, BAG3-IPV1, or BAG3-IPV2 mutants, or DsRed (Ctrl), demonstrating impairment in aggregation suppression by IPV mutations. Data presented as mean ± SEM from N = 3/4 biologically independent experiments and a total of n=17/12 replicated for IPVmut1/IPVmut2 respectively. (E) Representative confocal microscopy images cells expressing FUS-R521H-YFP (green), immunostained for LC3B (magenta), cell boundaries outlined in red. LC3B-positive puncta across shown as white dots. Scale bars - 10 µm for low magnification images and 5 µm for high magnification images. (F) Quantification of FUS aggregates colocalization with LC3B punctae. Cells expressing FUS-R521H-YFP were treated with MG-132 or DMSO (left), or transfected with siBAG3 or siControl (right). Cells were subsequently immunostained for LC3B, and colocalization of FUS aggregates with LC3B positive puncta was quantified from confocal images using Manders’ overlap coefficient, as described in the Methods. Bars represent the mean Manders’ overlap coefficient calculated from individual cells. Data are presented as mean ± SEM from N = 2 biologically independent experiments. For the MG-132 experiment, a total of n=212 DMSO treated and n=193 MG-132 treated cells were analyzed. For the siRNA experiment, n=532 siControl treated and n=518 siBAG3 treated cells were analyzed. (G) The autophagy inhibitor Bafilomycin A (200nM for 18h) was introduced in cells expressing FUS-R521H-YFP, co-transfected with either BAG3 or DsRed (Ctrl). Cells were subjected to PulSA analysis, showing that the suppression of FUS aggregation by BAG3 is severely impaired when blocking autophagy. Data presented as mean ± SEM from N=3 biologically independent experiments, with a total of n = 9 replicates per condition. (C, D, G) Dashed lines represent 95% CIs, ***p < 0.003, *p < 0.05, empirical p-value (see Methods). When two sample groups were compared to each other, rather than to the control, we report a two-sided t-test p-value.

To better understand the contribution of BAG3-HSP70 interactions to its inhibitory effect on mutFUS aggregation, we introduced a Q488A point mutation in the BAG domain (as described in ^37^). This mutant exhibited a significant reduction in HSP70 binding (Fig. 4B, S4A,B). Further testing of the functional effects of the Q488A mutation on mutFUS aggregation showed that BAG3-Q488A lost its ability to inhibit FUS-R521H-YFP aggregation (Fig. 4C). Thus, HSP70 binding to BAG3 is correlated with its function in suppressing mutFUS aggregation.

BAG3 was previously shown to enhance selective autophagy^33, 38, 39^. We thus next investigated the contribution of selective autophagy through BAG3 to mutFUS aggregation suppression. As the N-terminus of BAG3 contains a WW domain previously shown to be involved in BAG3-mediated autophagy^40^, we tested a WW-domain truncation mutant of BAG3 (ΔN, Fig. 4A). We found that indeed, in the absence of the WW domain, BAG3 lost its ability to inhibit mutFUS aggregation (Fig. 4C). ΔN-BAG3 exhibited a small, 15% reduction in HSP70 binding compared to the full length BAG3 (Fig. 4B, S4A,B), and therefore we could not exclude that loss of mutFUS aggregation suppression activity was confounded by reduced HSP70 recruitment. We therefore generated point mutations within the BAG3 IPV motifs (Fig. 4A), previously shown to mediate its interactions with sHSPs^39, 41^, which are important for autophagy-enhancing functions^33, 42^. Importantly, the interaction of these mutants with HSP70 was the same as that of the full length BAG3 (Fig. 4B, S4A,B), yet their ability to prevent mutFUS aggregation was significantly impaired (Fig. 4D).

Having established that BAG3 domains required for selective autophagy are also necessary for suppression of mutFUS aggregation, we next sought to directly assess the involvement of autophagy in this process. We therefore examined the co-localization of mutFUS aggregates with autophagosomes by immunofluorescence and quantitative image analysis, as a measure of autophagic targeting of mutFUS aggregates (see Methods). While FUS aggregates highly co-localized with LC3B punctae under basal conditions, their co-localization following MG-132 treatment indeed increased nearly two-fold (Fig. 4E,F). Interestingly, upon knockdown of BAG3, co-localization of FUS aggregates with LC3B punctae was significantly reduced (Fig. 4F), indicating that FUS aggregates are indeed targets for selective autophagy by BAG3. Lastly, we treated cells co-expressing mutFUS and BAG3 with the autophagy inhibitor Bafilomycin A, and analyzed mutFUS aggregation levels by FACS. The results showed that, in the presence of autophagy inhibitors, the ability of BAG3 to inhibit FUS aggregation was severely impaired (Fig. 4G). These data collectively suggest that BAG3 modulates mutFUS aggregation through activation of selective autophagy.

### SAT1 and the polyamine pathway as novel mediators of FUS aggregation suppression downstream of proteasome inhibitors

To identify additional mediators of the transcriptional program uniquely induced by proteasome inhibitors, we reanalyzed our RNA-seq data and systematically looked for genes that were uniquely upregulated by MG-132 treatments, while remaining completely unchanged under heat shock conditions (see Methods). This analysis identified 214 potential candidates that were induced only by MG-132 and not by HS (Fig. 5A, Table S1). Functional characterization of top hits using FACS found that SAT1, which showed a 25-fold induction by MG-132 treatment (Table S1), exhibited a substantial effect with approximately 80% mutFUS aggregation reduction (Fig. 5B, S5A, S1F).

**Figure 5:**
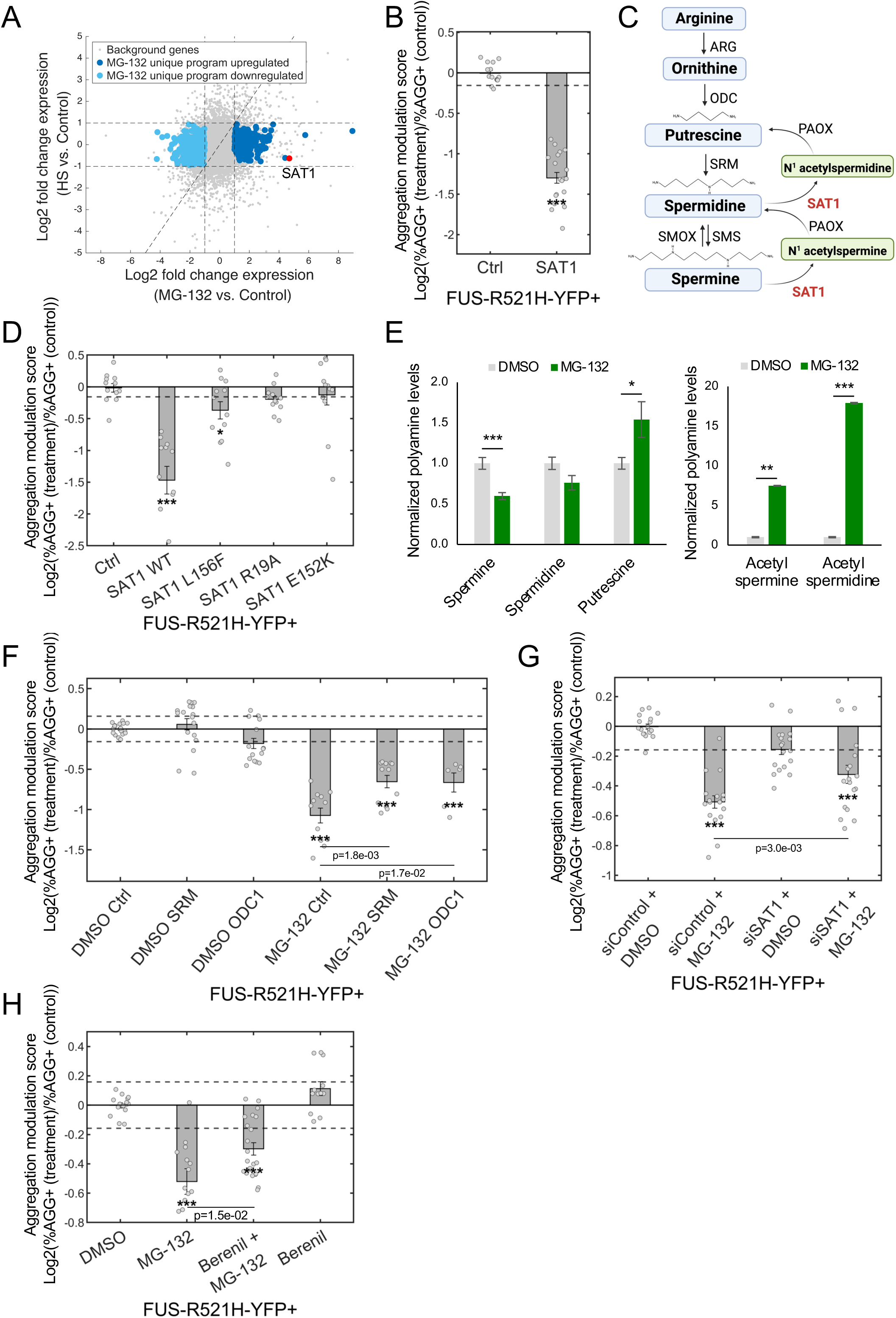
SAT1 and the polyamine pathway as novel mediators of FUS aggregation suppression downstream of proteasome inhibitors. (A) Log2 fold changes (LFC) of gene expression for cells treated with MG-132 (10uM for 18h) compared to control (x-axis) plotted against that of genes in cells subjected to heat shock (42°C for 8h) compared to their respective controls (y-axis). Dashed lines mark diagonal and LFCs of ± 1. Genes uniquely induced/repressed by MG-132 are marked in blue. SAT1 in red. (B) Aggregation modulation score for cells co-expressing FUS-R521H-YFP and SAT1 or DsRed (Ctrl) show major suppression of aggregation by SAT1. Data presented as mean ± SEM of N =4 biologically independent experiments, with a total of n =20 total replicates. (C) Schematic representation of the polyamine metabolic pathway. SRM, spermidine synthase; SMS, spermine synthase; ODC1, ornithine decarboxylase; ARG, arginases; SAT1, spermidine/spermine N1-acetyltransferase; PAOX, acetylpolyamine oxidase; SMOX, spermine oxidase. (D) FUS-R521H-YFP Aggregation modulation scores for SAT1 mutants, showing that metabolically inactive SAT1 loses its ability to inhibit FUS aggregation. Data presented as mean ± SEM of N =5 biologically independent experiments, with a total of n = 12 replicate. (E) Polyamine metabolomics demonstrates alterations in polyamine levels in cells treated with MG-132 compared to DMSO. Results were normalized to total protein measured by BCA. Data presented as mean ± SEM of N=3 independent biological experiments, with a total of n =8 replicates per condition. p=2.8e-4, 0.03, 1.1e-3, and 5.4e-4 for Spermine, Putrescine, Acetyl spermine and Acetyl spermidine respectively, calculated using t-test, two-sided. (F) Aggregation modulation scores for cells expressing FUS-R521H-YFP were co-transfected with SRM, ODC1, or DsRed (Ctrl) and treated with MG-132 or DMSO. Data are presented as mean ± SEM from at least N=3 biologically independent experiments, n = 6-16 total replicates per condition. (G) Aggregation modulation scores in cells expressing FUS-R521H-YFP that underwent SAT1 knockdown vs. siControl, and treated either with MG-132 or DMSO, show that SAT1 knockdown significantly reduced the suppressive effects of MG-132 on mutFUS aggregation. Data presented as mean± SEM of N = 6 biologically independent experiments, with a total of n = 14/18 replicates for siControl+DMSO/all other conditions, respectively. p-value calculated using paired t-test. (H) Aggregation modulation scores of cells expressing FUS-R521H-YFP, treated with either MG-132 (10uM, 18h), the SAT1 inhibitor Berenil (100uM, 18h), or both, further demonstrate that SAT1 inhibition impaired MG-132 suppressive function of FUS aggregation. Data are presented as mean ± SEM of N=5 independent biological experiments, with a total of n=12-22 replicates per condition. (B,D,F,G,H) Dashed lines represent 95% CI. ***p < 0.003, *p < 0.05, empirical p-value (see Methods). When two sample groups were compared to each other, rather than to the control, we report a two-sided t-test p-value.

SAT1, a polyamine N1-acetyltransferase, functions as the rate-limiting enzyme in polyamine catabolism and plays a central role in maintaining polyamine homeostasis^43^ (Fig. 5C). To further examine the potential interplay between the polyamine pathway and SAT1-mediated inhibition of mutFUS aggregation, we generated several SAT1 mutants with impaired metabolic activity: the E152K and R19A mutants previously reported to exhibit reduced binding to polyamines^44^, and the L156F mutant shown to have impaired enzymatic activity and low substrate binding^45^. PulSA analysis showed that, in the presence of all three metabolically impaired mutants, SAT1’s ability to inhibit mutFUS aggregation was abolished (Fig. 5D).

Next, we turned to explore potential connections between polyamine metabolism and proteasome inhibitors. To this end, we performed polyamine metabolomics (see Methods) to assess alterations in polyamine levels in cells treated with MG-132 compared to DMSO-treated controls. The levels of Spermine, a SAT1 substrate, were significantly decreased by 1.5-2 fold, while Putrescine showed a 1.5 increase following MG-132 treatment (Fig. 5E). Additionally, metabolomics demonstrated a marked accumulation in SAT1’s acetylated products, namely N1-Acetylspermidine (17.9 fold) and N1-Acetylspermine (7.5 fold), upon MG-132 treatment (Fig. 5E), indicating that proteasome inhibitors substantially alter polyamine metabolism. To further investigate the involvement of the polyamine pathway in FUS aggregation modulation, we examined two additional enzymes in the pathway, ODC1, and SRM, responsible for the production of Putrescine and Spermidine, respectively, and are therefore expected to show an opposite effect to that of the polyamine catabolic enzyme SAT1. While overexpression of each of these did not affect FUS aggregation in DMSO-treated cells, it partially impaired FUS aggregation suppression following MG-132 treatment (Fig. 5F), further implicating the polyamine pathway in FUS aggregation modulation.

To further establish SAT1 as a mediator of FUS aggregation suppression by proteasome inhibitors, we knocked down SAT1 in cells expressing FUS-R521H-YFP, and then treated them with MG-132 or DMSO. Introduction of siSAT1 into cells led to a reduction of 74-80% in its endogenous levels (Fig. S5B). MG-132 treatment in siSAT1 cells showed a significant impairment in aggregation suppression compared to siControl cells (Fig. 5G). Finally, we treated cells with Berenil (diminazene aceturate), which was shown to inhibit SAT1 activity^46^, either alone, or in combination with MG-132. Notably, Berenil partially but significantly hindered FUS aggregation suppression by MG-132 (Fig. 5H), further implicating SAT1’s central role in the effects of proteasome inhibitors on mutFUS aggregation.

### Proteasome inhibitors suppress FUS aggregation and alter polyamine metabolism in primary neurons

We next sought to examine the functional effects of proteasome inhibitors on mutFUS aggregation in primary neurons, using the BBB-permeable proteasome inhibitor Marizomib^25^. To this end, we used AAV-based viral infection to express FUS-R521H-YFP in primary rat cortical neuron cultures (Fig. S6A-F), as well as in primary mouse motor neuron (MN) cultures (Fig. S6B,I). Because FUS-R521H-YFP aggregates exhibit a complex morphology in neurons (Fig. S6B-D), and neuronal vulnerability precluded analysis by FACS, we used confocal microscopy images of live neurons followed by image analysis to quantify aggregation in neurons using a classifier trained to distinguish between nuclear fluorescent pattern and cells displaying the complex FUS-R521H-YFP aggregate phenotype, as in Rozales et al.^17^ (see Methods). We then calculated the percentage of aggregate-containing neurons out of all infected neurons, in FUS-R521H-YFP cortical or motor neuron cultures treated with Marizomib (Fig. 6A-C, S6F-I, S6K-M). We note that proteasome inhibitors did not significantly reduce FUS-R521H-YFP levels in neurons, as measured by western blot (Fig. S6J). Importantly, we found that proteasome inhibitors significantly reduced mutFUS aggregation in both cortical and motor neurons (Fig. 6A-C, S6F-I, S6K-M). FUS-R521H-YFP cortical neurons showed on average 27% less aggregate-containing neurons when treated with Marizomib compared to control (Fig. 6A,B, S6F,L, see Methods), and a 30% reduction was observed when applying MG-132 (Fig. S6G,H,K). Furthermore, in motor neurons, we found an even greater effect, with a 45% reduction in the percentage of aggregate-containing cells in response to Marizomib treatment (Fig. 6A,C, S6I,M).

**Figure 6:**
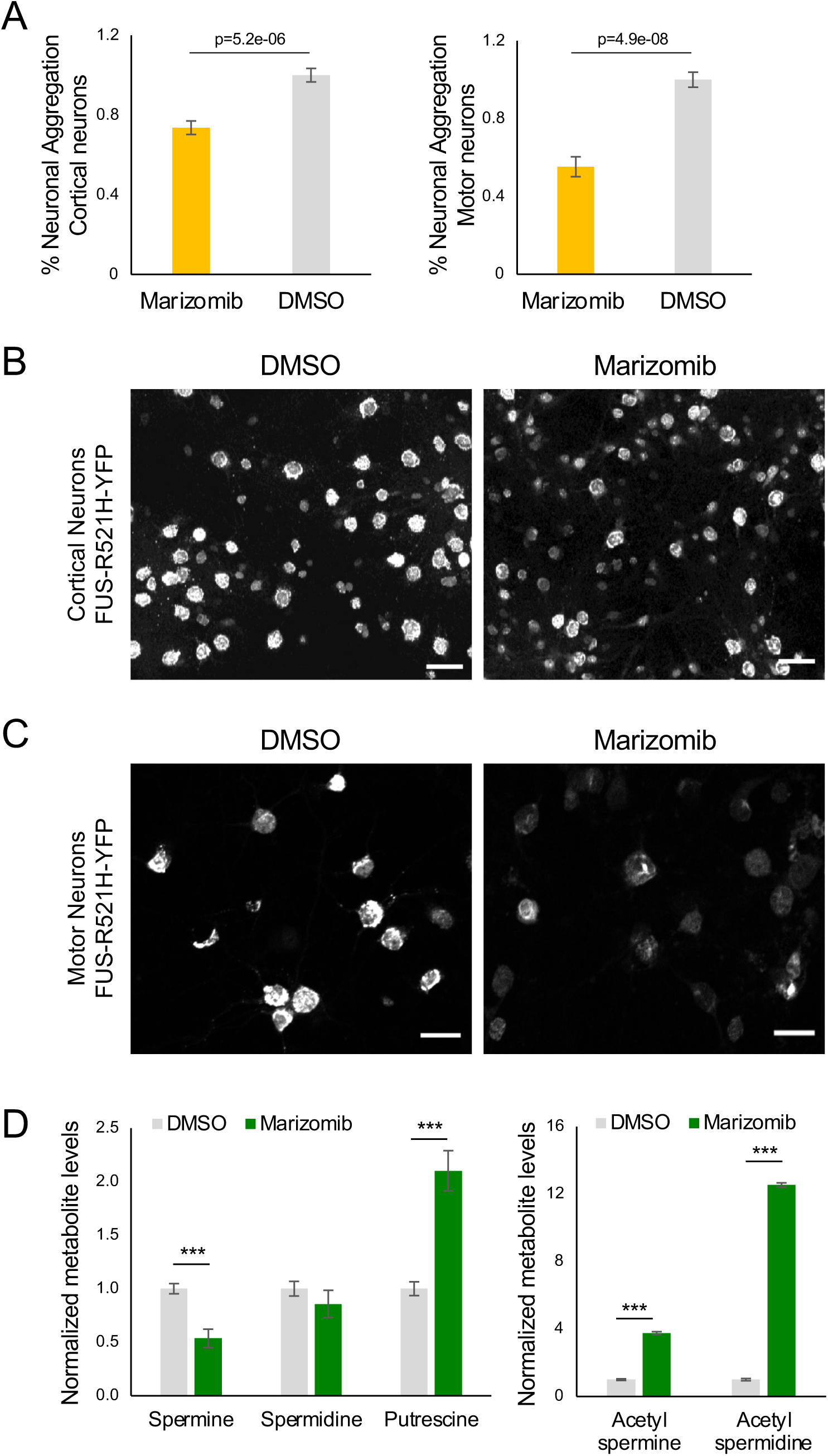
Proteasome inhibitors suppress FUS aggregation and affect polyamine metabolism in primary neurons. (A) Primary cortical neurons (left) or motor neurons (right) expressing FUS-R521H-YFP were treated with Marizomib (80 nM for cortical neurons or 60 nM for motor neurons) or DMSO as control. The fraction of aggregate positive neurons was quantified by live cell image analysis, and normalized to the mean of the corresponding DMSO condition within each biological replicate. Data presented as mean ± SEM from N = 3/2 biologically independent experiments for cortical/motor neurons. Quantification of individual experiments is shown in Fig. S6L,M. In total, n = 1,658/1,653 DMSO/Marizomib treated cortical neurons, and n = 567/362 DMSO/Marizomib treated motor neurons were analyzed across all experiments. Marizomib treatment significantly reduced the fraction of aggregate positive neurons in both cell types, p-values were calculated using a two-sided t-test. (B) Cortical neuronal cultures infected with FUS-R521H-YFP and treated with Marizomib (right panel) or DMSO as a control (left panel). Shown are maximum projection of confocal microscopy images (using Fiji), of representative fields out of N = 3 biologically independent experiments. Scale bar - 50 μm. Additional fields shown in Fig. S6F. (C) Motor neuron cultures infected with FUS-R521H-YFP and treated with Marizomib (right panel) or DMSO (left panel). Shown are sum projection of confocal microscopy fluorescent images (using Fiji), of representative fields out of N = 2 biologically independent samples. Scale bar - 24 μm. Additional fields shown in Fig. S6I. (D) Polyamine metabolomics was performed to assess alterations in polyamine levels in FUS-R521H-YFP expressing cortical neurons treated with Marizomib compared to control, results were normalized to total protein measurements by BCA. Shown are mean ± SEM of N=3 independent biological experiments, with a total of n=11/12 replicates for control/Marizomib respectively. p=1.5e-4, 2.6e-5, 3.9e-5, and 2.9e-6 for Spermine, Putrescine, Acetyl-spermine and Acetyl-spermidine respectively, calculated using t-test, two-sided.

At the molecular level, Marizomib treatment led to a significant induction in BAG3 levels in primary neurons (Fig. S6N), recapitulating the findings above (Fig. 3B). Finally, we asked whether the effects of proteasome inhibitors on the metabolism of polyamines that we observed in HEK293T cell models were also recapitulated in primary neurons. We treated primary cortical neurons with Marizomib and performed polyamine metabolomics. Indeed, mirroring the effects seen in cell models, we found significant changes in the levels of Spermine and Putrescine, while the levels of N1-Acetylspermidine and N1-Acetylspermine were markedly induced (Fig. 6D).

Collectively, our findings establish that proteasome inhibitors, including the BBB-permeable compound Marizomib, suppress mutFUS aggregation in both cell models and primary neurons, highlighting their therapeutic potential for FUS-ALS.

## Discussion

Here we revealed an unexpected suppressive effect of proteasome inhibitors on mutFUS aggregation, which include MG-132, Velcade (Bortezomib), and the BBB permeable proteasome inhibitor Marizomib. These findings were surprising, given the common notion that proteasome inhibitors lead to the accumulation of protein aggregates, due to impaired degradation of misfolded proteins by the UPS, as indeed known for many pathological aggregate types^10, 12, 23, 24^ (Fig. 1A,B).

The impact of proteasome inhibition on protein aggregation appears to be protein- and disease subtype-specific. Our results portray a model, in which proteasome inhibition simultaneously activates detrimental and protective proteostasis pathways, and the selective sensitivity of different aggregation-prone proteins to each of these determines whether aggregation increases or decreases (Fig. 7). On the one hand, inhibiting the UPS directly leads to the accumulation of misfolded proteins, exacerbating aggregation of various disease-associated drivers, such as HTT-polyQ, TDP-43^23, 24^ (Fig. 1A,B), and others^12^. On the other hand, proteasome inhibitors upregulate an HSF1-mediated proteostasis program, that can potentially promote the clearance of protein aggregates, through selective autophagy and other mechanisms (Fig. 7). Indeed, we found that FUS aggregates were strongly associated with autophagosomes, and this association increased upon proteasome inhibition (Fig. 4E,F). Curiously, while HTT-polyQ aggregates did not associate with autophagosomes under basal conditions, and TDP-43-ΔNLS-2KQ aggregates did co-localize with autophagosomes, the co-localization of both was significantly increased upon MG-132 treatment (Fig. S7A-C). Nonetheless, our data collectively shows that the combined effect downstream of proteasome inhibitors promotes the aggregation of both HTT-polyQ and TDP-43. FUS is therefore a unique case for which the beneficial arm downstream of proteasome inhibitors dominates, leading to aggregation inhibition.

**Figure 7:**
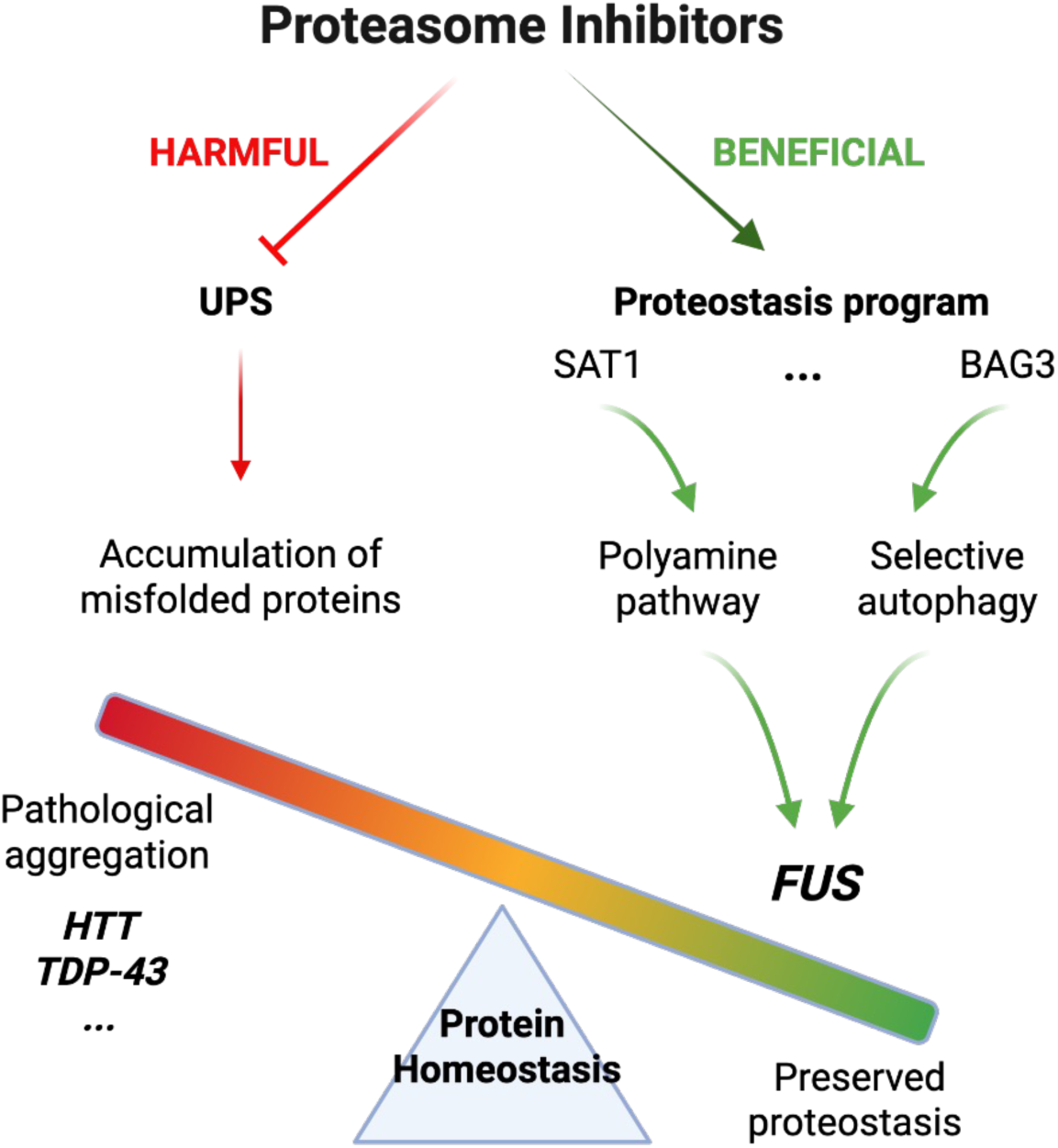
The dual effects of proteasome inhibition on protein aggregation and proteostasis. While inhibition of the ubiquitin-proteasome system (UPS) leads to the accumulation of misfolded proteins and worsens aggregation of various neurodegeneration-associated proteins, it concurrently activates proteostasis pathways that can fight aggregation. Among these, BAG3-mediated selective autophagy, and SAT1 induction, play critical roles in suppressing FUS aggregation. These opposing effects highlight the substrate-specific nature of proteasome inhibition effects on protein quality control mechanisms and on pathological aggregation.

Seeking to elucidate the underlying mechanism of this unique response of FUS to proteasome inhibitors, we found that the suppression of FUS aggregation by proteasome inhibitors was transcriptionally mediated, via HSF1 (Fig. 2). Indeed, it is known that proteasome inhibitors lead to a heat-shock response-like, yet not completely characterized, transcriptional response^47^. One key mediator that we identified as part of this program was BAG3, a co-chaperone of HSP70^35^, whose induction following proteasome inhibition was previously documented^10, 48, 49, 50^, and which was found to control the disassembly of stress granules containing WT FUS^51^. Prior studies indeed showed that the induction of BAG3 by MG-132 is regulated by HSF1^10, 52^. While BAG3 has been proposed to facilitate the autophagic clearance of protein aggregates associated with several neurodegenerative diseases^13, 39, 53, 54, 55^, suggesting a broader role in pathological aggregate clearance, our findings also highlight its substrate specificity. In agreement with a recent report^34^, we found that BAG3 enhanced, rather than suppressed, TDP-43 aggregation (Fig. S3D). Thus, unlike other aggregation-prone proteins, for which the detrimental effects of proteasome inhibition predominate, the selective activity of BAG3 toward FUS, together with its ability to promote selective autophagy, appears to outweigh the consequences of impaired proteasomal degradation, resulting in reduced FUS aggregation (Fig. 7).

Our data revealed the involvement of the polyamine pathway, and SAT1 in particular, in the suppression of FUS aggregation by proteasome inhibitors. Manipulating SAT1 and its enzymatic activity significantly altered mutFUS aggregation, identifying polyamine metabolism as a previously unrecognized regulator of FUS aggregation. Dysregulation of polyamine homeostasis has been implicated in several neurodegenerative diseases^56, 57, 58, 59^ and increased SAT1 activity was previously shown to alleviate α-synuclein toxicity^59^, suggesting that modulation of this pathway may represent a broader mechanism for combating pathological protein aggregation. Interestingly, spermidine has been reported to promote autophagy, suggesting a potential link between SAT1 and the BAG3-dependent selective autophagy pathway identified here. However, because spermidine levels remained unchanged following proteasome inhibition in both cell line models and primary neurons, our data suggest that the effects of SAT1 on FUS aggregation cannot be simply explained by altered spermidine abundance. Alternatively, polyamine metabolism may influence the proteotoxic stress response more directly, as polyamine depletion has been shown to impair HSF1 activation and inducible HSP70 expression^60^. The precise molecular connection between SAT1 activity, polyamine metabolism, HSF1 and FUS aggregation therefore remains to be elucidated.

Overall, our study highlights a unique proteostasis program downstream of proteasome inhibitors, involving BAG3-mediated selective autophagy, and polyamine metabolism alterations, which we revealed to be specifically beneficial towards battling FUS aggregation (Fig. 7). Importantly, although proteasome inhibitors were proposed as major drivers of TDP-43 aggregation^61^, in response to Marizomib treatment, WT TDP-43 formed transient nuclear condensates which completely dissolved after inhibitor removal (Fig. S7D). Since TDP-43 is not found to co-aggregate together with FUS in ALS^62, 63, 64^, our findings thus highlight BBB-permeable proteasome inhibitors, such as Marizomib, as potential novel avenues for therapeutic intervention specifically for FUS-ALS.

## Methods

### Cloning and plasmid preparation

pTRex-FUS-WT-YFP, pTRex-FUS-R521H-YFP, and pTRex-FUS-R518K-YFP, pTRex-FUS-R521C-YFP, pTRex-FUS-R495X-YFP, pCAG-FUS-R521H-YFP, pTRex-HTT-134Q-GFP, and pcDNA3-BAG3-3xFLAG-V5 constructs were described in Rozales et al. ^17^. Human TDP-43 was cloned into pENTR221-YFP, and site-directed mutagenesis was performed using Quikchange 2 Site-Directed Mutagenesis Kit (Agilent Technologies 210518), to create 2 K→Q mutation at residues K145 and K192, in addition to mutations in the nuclear localization sequence (NLS): K82A, R83A, K84A to generate TDP-43-ΔNLS-2KQ^29^. This clone was then transferred to a pAAV-ccdb backbone using gateway cloning. SAT1 (WT) was cloned from HEK293T cDNA using PCR (see primers Table S2) into a pcDNA3.1-ccdb-3xFLAG-V5 plasmid using gateway cloning. The different point mutants for BAG3 and SAT1 were generated using QuikChange2 Kit (see primers in Table S2). The BAG3-ΔC isoform and the ΔN-BAG3 were both generated from the full-length isoform (WT isoform) using PCR and gateway cloning, (see primers in Table S2). The polyamine pathway genes SRM and ODC1 were obtained from the human ORFeome collection (GE Healthcare) and subcloned from pDONR223 into the pcDNA3.1-ccdB-3xFLAG-V5 expression vector using gateway cloning. All plasmid sequences are provided in Table S3.

### Cell culture, transfections and siRNA

HEK293T cells were grown in standard DMEM supplemented with 10% FBS, 1% Penicillin-Streptomycin in 37 °C, 5% CO_2_ incubator. For the FUS transfections, HEK293T cells (3e5 cells per well) were seeded on 6well plates and transfected using PEI transfection reagent (Thermo Fisher Scientific) one day after seeding, according to the manufacturer instruction, with 1250ng of FUS DNA and 1250ng of chaperone DNA/DsRed as control. Cells were assayed 48 h following the transfection. For siRNA experiments, cells were seeded at 150,000 cells per 6well plate one day before transfection. siRNA transfections were performed using RNAiMAX lipofactamine reagent (Thermo Fisher Scientific), according to the manufacturer’s protocol, with siRNA concentration of 15nM (Dharmacon Smartpool, L-012109-02-0005 for siHSF1, L-011957-00-0005 for siBAG3 and L-010382-00-0005 for siSAT1) or siControl (Dharmacon Smartpool, 001810-10-05 or 020585-01-0005). One day after transfection of the siRNA cells were co-transfected with 1250ng of FUS-R521H and 1250ng of DsRed using PEI. Cells were analyzed 48 h later.

### PulSA analysis, Aggregation Modulation score and p-value

For PulSA analysis, cells were collected 48 h after transfection, cells were trypsinized, resuspended in media with serum, placed on ice, and DAPI regent was added (5 µM) in order to assay cell viability. Cells were then taken to Flow cytometry in order to differentiate between subpopulations with diffused cellular fluorescence and those with protein aggregates^28^, as in Rozales et al. ^17^ (Fig. S1A). To obtain Aggregation modulation scores, cells of appropriate size were considered during the FACS analysis, and non-viable cells (positive for DAPI) were excluded. We then quantified the percentage of cells that showed a fluorescent aggregate pattern (AGG+), and compared these percentages between treated and untreated cells, or cells co-transfected with a modulators compared to cells co-transfected with DsRed as a control. Aggregation modulation scores for each modulator or treatment were calculated as log2(%AGG+ (modulator or treatment)/%AGG+ (control)). Confidence intervals (95%, appear as dashed lines) were calculated according to the variations between different DsRed co-transfected cells replicates, such that they represent twice the standard deviation (2*STD) of Aggregation modulation scores calculated between many replicates of the control, as described in Rozales et al.^17^. Chaperones or treatments were called as significant modulators if their mean Aggregation modulation scores +/− SEM was below/above the 95% CI in the case of a negative/positive score respectively. An empirical *p*-value was indicated as *p* < 0.05 (*) if the score value was below/above the 95% CI (i.e. twice the STD), and *p* < 0.003 (***) if the score was below the 99.7% CI (i.e., 3 times STD). HTT CI = +/-0.1168 based on n=57 replicates, FUS CI = +/- 0.1577 based on n=90 replicates, as previously determined^17^. TDP-43 CI = +/-0.1612 was determined in this study based on n=28 replicates. This empirical method thus incorporates both statistical confidence and requirement on effect size relative to control variability. In addition, whenever two sample groups were compared to each other, rather than to the control, we report a two tailed t-test p-value.

Flow cytometry gates for AGG+, AGG-, and non transfected cell populations were maintained constant throughout the study, after being initially chosen to encompass the three populations that naturally appeared in the data (using contour plots, Fig. S1B). Nonetheless, all the results were robust to the choice of FACS gating strategy, as varying the gates maintained significant trends in all experiments, and the choice of gating used to generate scores throughout this study was the most strict (Fig. S1F).

FUS-R521H-YFP aggregation suppression by proteasome inhibitors is robust to FUS-R521H-YFP expression levels (Fig. 1G, S1K). We verified that aggregation suppression is observed when controlling for expression levels between the AGG+ and AGG-populations. To this end, we divided cells by FUS-R521H-YFP levels, using flow cytometry GFP-H cell values, into bins defined as equal quantiles of the log10(GFP-H) value distribution across the AGG-population. We then plotted on the y-axis, for each expression bin, the percentage of AGG+ cells out of all cells, in the DMSO control (grey) or the proteasome inhibitor-treated (red) sample, similarly to^65^. This analysis demonstrated that, for both MG-132 and Marizomib, the percentage of AGG+ cells was consistently lower than the percentage in the DMSO control across all expression bins.

### RNA-seq and data analyses

Cells were subjected to the following treatments: MG-132 for 7 and 18 hours with DMSO as a control, with N=2 biological replicates, and HS at 44°C for 2h or 42°C for 8h, with no treatment as control, with N=4/4/3 replicates, respectively. PolyA selected, strand specific RNA-seq was performed as described in Rozales et al. ^17^. Fold change was calculated using Deseq2 ^66^. In Fig. 5A, in grey, background, all expressed genes (TPM >=1). In blue, MG-132 unique program, defined as genes that were significantly changed after MG-132 treatment at both 7h and 18h (Deseq2 |LFC|>1, Deseq2 padj<0.01) and were not significantly changing after HS treatment both at 2h and 8h treatments (Deseq2 |LFC|<1, Deseq2-padj>0.05). See Table S1 for gene lists. RNA-seq presented in Fig. 3B shows a scatter plot of log2 fold change of MG-132 18h vs. control (x-axis), compared to heat shock (42°C) 8h vs. control (y-axis), for the indicated functional groups only. Chaperones families and functional groups in Fig. 3A,B were color coded, chaperone family assignments were taken form Brehme et al.^67^, ubiquitin ligases were downloaded from Uniprot, autophagy core and regulators lists were taken from^68^.

### Immunofluorescence staining

HEK293T cells were cultured on coverslips and transfected as above. Two days post transfection, cells were fixed in 4% paraformaldehyde (PFA) in PBS, permeabilized with 0.5% Triton-x100 in IF buffer (5% FCS, 2% BSA in PBS), then immunostained by primary anti-FLAG antibody (Sigma-Aldrich, F1804) followed by secondary AlexaFluor 647 Donkey Anti-Mouse (Jackson ImmunoResearch) antibody to image FLAG-tagged chaperones. The far-red secondary antibody was used to eliminate fluorescence overlap between the YFP/GFP-tagged aggregates and the chaperones. Cells were additionally stained with DAPI to mark nuclei.

For immunostaining of primary cortical neurons, neurons grown on coverslips were washed with Tyrode’s physiological solution, fixed using 4% PFA and 4% sucrose for 20 min at room temperature, and permeabilized with 0.25% Triton X-100 for 20 min. Following blocking in immunofluorescence (IF) buffer (as described above) for 30 min, cells were incubated overnight at 4 °C with primary antibodies diluted in IF buffer, including mouse monoclonal anti-FUS/TLS (Santa Cruz Biotechnology, sc-47711, 1:200) and/or rabbit monoclonal anti-MAP2 (Cell Signaling Technology, 4542S, 1:400). After washing, cells were incubated overnight at 4 °C with Alexa Fluor 488-conjugated secondary antibody (1:400) and Cy5-conjugated goat anti-rabbit secondary antibody (Jackson ImmunoResearch; 1:400). Nuclei were stained with DAPI (1 μg/mL) for 5 min at room temperature. Coverslips were mounted using DAKO mounting medium and imaged by confocal microscopy.

### Aggregation Quantification by fluorescence microscopy Imaging

Nuclei were segmented using Cellpose^69^. Protein aggregates were detected in the fluorescence channel of each protein (FUS-R521H–YFP, HTT-polyQ–GFP, TDP-43-△NLS-2KQ-YFP) by Difference-of-Gaussians background subtraction followed by Otsu thresholding on the maximum-intensity projection. Aggregates outside a size range of 2.2–128 µm² were excluded, and each aggregate was assigned to its nearest nucleus. Cells were classified as AGG+ (one or more aggregates), AGG− (no aggregates, GFP intensity above expression threshold), or GFP− (non-expressing). Expression thresholds were applied to the diffused signal intensity for each protein. Analyses were performed using a custom Python pipeline.

### LC3B Immunofluorescence and Aggregate colocalization Analysis

Cells expressing FUS-R521H–YFP, HTT-polyQ–GFP, or TDP-43-△NLS-2KQ-YFP were treated with MG-132 (10 µM, 18 h) or DMSO as a control, or transfected with siBAG3 or siControl for FUS samples only (see Methods). Cells were methanol fixed and immunostained for LC3B (Cell Signaling Technology, C3868), followed by Cy5 647 Goat anti Rabbit (Jackson ImmunoResearch) to visualize autophagosomes. The immunostaining procedure proceeded as in the Immunofluorescence Methods section. Images were acquired using a laser scanning confocal microscope (LSM700) with identical acquisition settings across each experimental set for the LC3B channel. Image analysis was performed using custom Python scripts. Cell instance segmentation was carried out using Cellpose^69^, enabling automated and low bias identification of individual cells. LC3B positive puncta were segmented by detecting blob like features using a Difference of Gaussian (DoG) based approach (skimage.feature.blob_dog). Following blob detection, a dataset specific threshold was applied to retain bona fide puncta and exclude spurious detections. FUS-LC3B colocalization was quantified using Manders’ coefficient, defined as the area of overlapping positive signals (the aggregates channel was thresholded using Otsu segmentation, whereas the LC3B channel threshold was defined by the minimum intensity of the puncta detected in the preceding step) divided by the total area of the aggregate-positive signal. Total cellular LC3B levels were quantified as the raw integrated LC3B fluorescence per cell, normalized to cell area.

### Western blotting and Co-immunoprecipitation

Cells were lysed in RIPA buffer ((50 mM Tris-HCl, pH 7.4, 150 mM NaCl, 1% NP-40, 0.5% sodium deoxycholate, 0.1% SDS) supplemented with protease inhibitors, followed by sonication (5 cycles of 5 seconds each) to ensure complete lysis. The lysates were then passed through a 21-gauge needle to further homogenize the samples. Protein concentrations were determined using the BCA assay (Thermo Fisher). Equal amounts of protein were loaded on the gels. Membranes were probed with the following primary antibodies: β-Actin (0869100-CF, MP Bio), and anti-GFP (MBL, #598), anti-FLAG (Sigma-Aldrich, F1804), anti-HSC70/HSP70 (Enzo Life Sciences, N27F34), GAPDH (Abcam, ab8245).

For co-immunoprecipitation (co-IP) of FLAG-tagged chaperones, lysis buffer was prepared as follows: 50 mM HEPES pH 7.9, 0.5% Triton X-100, 5% glycerol, 150 mM NaCl and 2 mM EDTA and Protease inhibitors (Sigma-Aldrich P5726). Wash buffer was prepared as follows: 50 mM HEPES pH 7.9, 1% Triton X-100, 5% glycerol, 150 mM NaCl and 2 mM EDTA. Cells were transfected as above. At 48 h post transfection, cells were placed on ice, washed with ice-cold PBS, lysis buffer was added and cells were collected by scraping. Lysates were centrifuged (8200 g for 10 min at 4°C), and supernatant was transferred to new tubes. Samples were IP-ed using 10 µl of EZview Red anti-FLAG M2 Affinity Gel beads (Sigma-Aldrich, F2426), which were added to equal sample volumes of 600 µl for each sample and samples with beads were slowly rotated 3h at 4°C. Beads were then washed 7 times with 1 ml of wash buffer. Following the last wash, the beads were resuspended in equal volumes of wash buffer, protein sample buffer was added, and the samples were boiled at 100°C for 5 min. Equal volumes were then run on an 8% or a 12.5% gel, and immunoblotted using anti-FLAG (Sigma-Aldrich, F1804), anti-HSC70/HSP70 (Enzo Life Sciences, N27F34).

### Proteasome inhibitors and other drug treatments

Cells were seeded in 6-well plates at a density of 3.5 × 10⁵ cells per well and transfected as described above. Thirty hours after FUS transfection, cells were treated with the indicated compounds for 18 h unless otherwise specified. For HEK293T cells, MG-132 (Sigma-Aldrich, M7449) was used at a final concentration of 10 μM, Marizomib (Sigma-Aldrich, SML1916) at 80 nM (unless otherwise indicated), Velcade (Bortezomib; Selleck Chemicals, S1013) at 10 μM, and Withaferin A (Sigma-Aldrich, W4394) at 6 μM. Bafilomycin A (Sigma-Aldrich, B1793) was used at 200 nM for 18 h, and Berenil (diminazene aceturate; Sigma-Aldrich, D7770) was used at 100 μM for 18 h. DMSO was used as the vehicle control, with its final concentration matched to that of the corresponding treatment, which did not exceed 1:1000 in any of the treatments. Cells were analyzed 48 h after transfection.

For primary cortical neurons, MG-132 was used at 7.5 μM and Marizomib at 80 nM. For primary motor neurons, Marizomib was used at 60 nM.

### Polyamine Metabolomics analysis

HEK293T were seeded and transfected in 6-well plate, neurons were grown on 24-well plates coated with poly-D-Lysine. The plates were placed on ice, and the medium was discarded. The cells were then washed once with PBS at room temperature and ice-cold (−20°C) metabolite extraction solution (MeOH:ACN:Water; 5:3:2; v/v/v) was added to each well. The plates were shaken in at 4 °C for 20 minutes to complete the metabolite extraction process. After extraction, every three or four wells were combined (for HEK293T or neurons respectively) into one sample. The combined samples were centrifuged at 4°C for 15 minutes at maximum speed. The supernatant was transferred into a clean Eppendorf tube, dried down, and re-suspended in extraction solution. The samples were stored at −80°C before LC/MS. The samples were run in SIM mode to increase sensitivity. Analysis method: LC-MS metabolomics analysis was performed using DionexTM ultra high-performance liquid chromatography (UPLC) system coupled to Q-ExactiveTM Orbitrap Mass Spectrometer (ThermoFisher Scientific), with a resolution of 70,000 at 200 mass/charge ratio (m/z), in electrospray ionization at positive mode (3500 v), used at tSIM mode for spermine, spermidine, putrescine and their acetylated forms to increase sensitivity (e.g. the following m/z were followed). AGC target was set to 5E04 with max IT of 200 ms. RF level was set to 50, Sheath and Aux gas was set to 30 and 13 respectively (arbitraty units) with capillary tube temperature at 263 °C and Aux gas heater temperature at 425 °C. HPLC setup consisted of SeQuant® ZIC®-cHILIC TM (100 mm x 2.1 mm, 3µm. 10 µL of biological extracts were injected and the compounds were separated with mobile phase consisting of water with 0.1% formic acid (mobile phase A) of mobile phase B composed of Acetonitrile. Flow rate and column temperature were maintained at 0.4 mL/min and 45 °C, respectively, for a total run time of 16 minutes. Starting at 95% mobile phase B with a gradient lasting 10 minutes until reaching 90% mobile phase A. Polyamines and their acetylated forms were detected using a mass accuracy below 5 ppm. Xcalibur (ThermoFisher Scientific) was used for data acquisition. Data Analysis was performed using Skyline Daily^246^ where mass accuracy and chromatographic retention time were used for identification and peak integration using an in-house authentic standards library. Peak intensities were normalized to ug protein, measured a BCA kit.

### Neuronal cultures and infections

Viruses were packaged with the AAV-PHP.eB capsid as described in Rozales et al. ^17^. All viruses were quantified for titer using qPCR within the same assay, ensuring consistent infection titers across experiments. Isolation and culturing of rat primary cortical neurons were done as previously described in Rozales et al. ^17^ (Ethics certificate No. IL-130-09-17). Briefly, P0 neonates, from timed-pregnant white coat female Sprague Dawley female rats (Charles River; SAS SD, strain code 400), were euthanized and their cortex was dissected and dissociated. Then, 150 K cells were plated in 24-well flat bottom plates on coverslips (12 mm, thickness ∼0.1 mm) coated with Poly-D-Lysine (Sigma-Aldrich P7886-50MGP) in minimum essential medium (MEM), glutamine-free based Neuronal growth medium (5% fetal bovine serum, 2% B27, 1% Glutamax (100×, Invitrogen), 2% 1 mol/L d-glucose and 0.1% μL serum extender (BD Biosciences, Cat. No. 355006)). Cultures were incubated at 37 °C and 5% CO_2_. At 5 days after plating, media was changed and supplemented with cytosine arabinosine (Ara-C, Sigma, C6645) at a final concentration of 4 μM, to reduce the growth of glia in the cultures. Cultures were infected at day 5 with pAAV-FUS-R521H-YFP. After that, culture media was replaced every 48 hours, and cells were imaged 6 days post infection (see Image Analysis section below).

Primary spinal cord motor neurons were isolated from E12.5 mouse embryos and cultured as described in Zahavi et al., 2015^70^. Briefly, spinal cords were dissected, trypsinized, and mechanically dissociated by trituration. The resulting cell suspension was centrifuged through a 4% BSA cushion, after which the pellet was resuspended and purified by centrifugation through an OptiPrep gradient (10.4% OptiPrep; Sigma-Aldrich, 10 mM Tricine, 4% glucose) for 20 min at 760 × g with the brake turned off. Cells were collected from the interface, washed once with complete medium, and plated onto coated culture chambers. Neurons were maintained in Neurobasal medium (Gibco) supplemented with B27 (Gibco), 10% (v/v) horse serum (Gibco), 25 μM β-mercaptoethanol, 1% penicillin–streptomycin (Lifegene), 1% GlutaMAX (Gibco), 1 ng/mL GDNF, 0.5 ng/mL CNTF, and 1 ng/mL BDNF (Alomone Labs). Prior to plating, culture chambers were coated overnight at 37 °C with 1.5 μg/mL poly-DL-ornithine (Sigma-Aldrich), followed by 3 μg/mL laminin (Sigma-Aldrich) for 2 h at 37 °C. Cultures were then incubated at 37 °C and 5% CO_2_. Cultures were infected at day 1 post plating with pAAV-FUS-R521H-YFP. After that culture media was replaced every 48 hours. At 5 days after plating, media was changed and supplemented with cytosine arabinosine (Ara-C, Sigma, C6645) at a final concentration of 1 μM, to reduce the growth of glia in the cultures. Cells were imaged 6 days post infection (see Image Analysis section below).

### Neuronal Aggregation Image Analysis

For microscopy analysis, primary cultures were infected as above, and cells were then treated with Marizomib (80 nM for cortical neurons or 60 nM for motor neurons) or MG-132 (7.5 µM) for 18 hours.

Images were acquired with a Zeiss LSM 880 confocal microscope using water dipping Plan-Apochromat ×20 objective, (working distance, 1.8 mm). The acquired images were processed and analyzed using a Fiji script, as in Rozales et al.^17^. Out of each confocal microscopy image, fixed number of z-stacks were successively chosen along the z-axis and stacked to form one image. An equal number of regions of interest (651 × 651 μm in size) were chosen as input for the script. Then, total cell numbers per processed image were identified based on YFP intensity threshold (>240). Subsequently, separation of cells was performed based on watershed processing, cell filtering was based on size (>50 μm2), and cellular aggregate detection was performed using Trainable Weka Segmentation plugin, summing aggregate area per cell, where a total area >5 μm were considered aggregates. Finally, a Python script was used to extract output excel files.

The image-analysis classifier exhibited a low background error rate, as validated on images of FUS-WT-YFP–expressing neurons, consistent with previous findings (Rozlaes et al.^17^). In the MG-132 experiments, 741, 957, and 555 cells were analyzed in the respective control groups, and 831, 1185, and 278 cells in the corresponding MG-132 treated groups across three independent biological replicates. In the Marizomib experiments (cortical neurons), 1,077, 411, and 156 neurons were analyzed following Marizomib treatment, with 801, 709, and 148 neurons serving as matched controls across independent experiments. For motor neurons, 78 and 284 cells were analyzed in the Marizomib-treated groups, compared with 211 and 356 cells in the corresponding control groups across independent biological replicates.

Following image analysis, the number of aggregate containing cells was divided by the total number of fluorescent cells detected by the classifier to obtain a fraction of aggregate containing neuron for each image. Then, average and SEM for the fraction of aggregate containing neurons was calculated for each treatment vs. control in each experiment, and a two tailed t-test p-value was calculated accordingly.

To generate the averaged plot for all experiments (Fig. 6A, S6G), the mean aggregate fraction of the DMSO treated control was calculated separately for each biological replicate. Then, the aggregate fraction measured in each field was divided by the corresponding experiment-specific DMSO mean. The normalized values were subsequently averaged within each biological experiment across all replicates, and the final mean (and SEM) across all fields from all replicates was plotted.

## Acknowledgements

This work was supported by a grant from the Congressionally Directed Medical Research Programs (CDMRP), ALS-TIA, grant number HT9425231031. We thank the Prince Center for Neurodegenerative disorders of the brain, and the Rappaport Family Institute for Research in the Medical Sciences for their funding support. We thank the Technion Rappaport Faculty of Medicine Biomedical Core Facilities for their support in microscopy and in particular to Raghd Abu-Sinni for image analysis of autophagy.

## Author contributions

The project was conceived by RS. RS, AY, KY and KR designed all experiments. AY, KY and KR performed all experiments, they were assisted by all other authors. AM and NS analyzed RNA-seq data. RS and AY wrote the manuscript.

## Data Availability

The datasets generated during the current study are included in this published article and its supplementary information files. Raw data used to generate Aggregation Modulation Scores are available from the corresponding author upon request. RNA-seq data was deposited in GEO, GSE315527.

## References

1. Gitler AD, Dhillon P, Shorter J. Neurodegenerative disease: models, mechanisms, and a new hope. Disease models & mechanisms 10, 499–502 (2017).

2. Muchowski PJ, Wacker JL. Modulation of neurodegeneration by molecular chaperones. Nature Reviews Neuroscience 6, 11–22 (2005).

3. Kabashi E, et al. TARDBP mutations in individuals with sporadic and familial amyotrophic lateral sclerosis. Nat Genet 40, 572–574 (2008).

4. Kwiatkowski TJ, Jr., et al. Mutations in the FUS/TLS gene on chromosome 16 cause familial amyotrophic lateral sclerosis. Science 323, 1205–1208 (2009).

5. Sreedharan J, et al. TDP-43 mutations in familial and sporadic amyotrophic lateral sclerosis. Science 319, 1668–1672 (2008).

6. Taylor JP, Brown RH, Jr., Cleveland DW. Decoding ALS: from genes to mechanism. Nature 539, 197–206 (2016).

7. Hartl FU, Hayer-Hartl M. Molecular Chaperones in the Cytosol: from Nascent Chain to Folded Protein. Science 295, 1852–1858 (2002).

8. Labbadia J, Morimoto RI. The biology of proteostasis in aging and disease. Annu Rev Biochem 84, 435–464 (2015).

9. Bukau B, Weissman J, Horwich A. Molecular chaperones and protein quality control. Cell 125, 443–451 (2006).

10. Dikic I. Proteasomal and Autophagic Degradation Systems. Annu Rev Biochem 86, 193–224 (2017).

11. Bercovich B, et al. Ubiquitin-dependent degradation of certain protein substrates in vitro requires the molecular chaperone Hsc70. The Journal of biological chemistry 272, 9002–9010 (1997).

12. Ciechanover A, Kwon YT. Degradation of misfolded proteins in neurodegenerative diseases: therapeutic targets and strategies. Exp Mol Med 47, e147 (2015).

13. Crippa V, et al. The small heat shock protein B8 (HspB8) promotes autophagic removal of misfolded proteins involved in amyotrophic lateral sclerosis (ALS). Hum Mol Genet 19, 3440–3456 (2010).

14. Lackie RE, et al. The Hsp70/Hsp90 Chaperone Machinery in Neurodegenerative Diseases. Front Neurosci 11, 254 (2017).

15. Yu H, et al. HSP70 chaperones RNA-free TDP-43 into anisotropic intranuclear liquid spherical shells. Science 371, (2021).

16. Lu S, et al. Heat-shock chaperone HSPB1 regulates cytoplasmic TDP-43 phase separation and liquid-to-gel transition. Nat Cell Biol 24, 1378–1393 (2022).

17. Rozales K, et al. Differential roles for DNAJ isoforms in HTT-polyQ and FUS aggregation modulation revealed by chaperone screens. Nat Commun 13, 516 (2022).

18. Akaree N, Secco V, Levy-Adam F, Younis A, Carra S, Shalgi R. Regulation of physiological and pathological condensates by molecular chaperones. FEBS J, (2025).

19. Benatar M, et al. Randomized, double-blind, placebo-controlled trial of arimoclomol in rapidly progressive SOD1 ALS. Neurology 90, e565–e574 (2018).

20. Kieran D, Kalmar B, Dick JR, Riddoch-Contreras J, Burnstock G, Greensmith L. Treatment with arimoclomol, a coinducer of heat shock proteins, delays disease progression in ALS mice. Nat Med 10, 402–405 (2004).

21. Cudkowicz ME. HEALEY ALS Platform Trial (Master Protocol). (ed(eds) (2020).

22. Benatar M, et al. Safety and efficacy of arimoclomol in patients with early amyotrophic lateral sclerosis (ORARIALS-01): a randomised, double-blind, placebo-controlled, multicentre, phase 3 trial. The Lancet Neurology 23, 687–699 (2024).

23. Li X, et al. Inhibiting the ubiquitin-proteasome system leads to preferential accumulation of toxic N-terminal mutant huntingtin fragments. Hum Mol Genet 19, 2445–2455 (2010).

24. Scotter EL, et al. Differential roles of the ubiquitin proteasome system and autophagy in the clearance of soluble and aggregated TDP-43 species. J Cell Sci 127, 1263–1278 (2014).

25. Di K, et al. Marizomib activity as a single agent in malignant gliomas: ability to cross the blood-brain barrier. Neuro Oncol 18, 840–848 (2016).

26. Lee DH, Goldberg AL. Proteasome inhibitors: valuable new tools for cell biologists. Trends Cell Biol 8, 397–403 (1998).

27. Dantuma NP, Bott LC. The ubiquitin-proteasome system in neurodegenerative diseases: precipitating factor, yet part of the solution. Front Mol Neurosci 7, 70 (2014).

28. Ramdzan YM, et al. Tracking protein aggregation and mislocalization in cells with flow cytometry. Nat Methods 9, 467–470 (2012).

29. Cohen TJ, Hwang AW, Restrepo CR, Yuan CX, Trojanowski JQ, Lee VM. An acetylation switch controls TDP-43 function and aggregation propensity. Nat Commun 6, 5845 (2015).

30. Santagata S, et al. Using the heat-shock response to discover anticancer compounds that target protein homeostasis. ACS Chem Biol 7, 340–349 (2012).

31. Akerfelt M, Morimoto RI, Sistonen L. Heat shock factors: integrators of cell stress, development and lifespan. Nat Rev Mol Cell Biol 11, 545–555 (2010).

32. Mathew A, Mathur SK, Jolly C, Fox SG, Kim S, Morimoto RI. Stress-specific activation and repression of heat shock factors 1 and 2. Mol Cell Biol 21, 7163–7171 (2001).

33. Behl C. BAG3 and friends: co-chaperones in selective autophagy during aging and disease. Autophagy 7, 795–798 (2011).

34. Lu S, et al. TDP-43 skein-like inclusions are formed by BAG3- and HSP70-guided co-aggregation with actin-binding proteins. Nat Cell Biol 27, 1925–1937 (2025).

35. Bracher A, Verghese J. The nucleotide exchange factors of Hsp70 molecular chaperones. Front Mol Biosci 2, 10 (2015).

36. Rosenzweig R, Nillegoda NB, Mayer MP, Bukau B. The Hsp70 chaperone network. Nat Rev Mol Cell Biol 20, 665–680 (2019).

37. Gentilella A, Khalili K. BAG3 expression in glioblastoma cells promotes accumulation of ubiquitinated clients in an Hsp70-dependent manner. J Biol Chem 286, 9205–9215 (2011).

38. Gamerdinger M, Hajieva P, Kaya AM, Wolfrum U, Hartl FU, Behl C. Protein quality control during aging involves recruitment of the macroautophagy pathway by BAG3. EMBO J 28, 889–901 (2009).

39. Carra S, Seguin SJ, Lambert H, Landry J. HspB8 chaperone activity toward poly(Q)-containing proteins depends on its association with Bag3, a stimulator of macroautophagy. J Biol Chem 283, 1437–1444 (2008).

40. Merabova N, et al. WW domain of BAG3 is required for the induction of autophagy in glioma cells. J Cell Physiol 230, 831–841 (2015).

41. Fuchs M, et al. Identification of the key structural motifs involved in HspB8/HspB6-Bag3 interaction. Biochem J 425, 245–255 (2009).

42. Myers VD, et al. The Multifunctional Protein BAG3: A Novel Therapeutic Target in Cardiovascular Disease. JACC Basic Transl Sci 3, 122–131 (2018).

43. Pegg AE. Spermidine/spermine-N(1)-acetyltransferase: a key metabolic regulator. Am J Physiol Endocrinol Metab 294, E995–1010 (2008).

44. Coleman CS, Pegg AE. Proteasomal degradation of spermidine/spermine N1-acetyltransferase requires the carboxyl-terminal glutamic acid residues. J Biol Chem 272, 12164–12169 (1997).

45. McCloskey DE, Pegg AE. Properties of the spermidine/spermine N1-acetyltransferase mutant L156F that decreases cellular sensitivity to the polyamine analogue N1, N11-bis(ethyl)norspermine. J Biol Chem 278, 13881-13887 (2003).

46. Libby PR, Porter CW. Inhibition of enzymes of polyamine back-conversion by pentamidine and berenil. Biochem Pharmacol 44, 830–832 (1992).

47. Bush KT, Goldberg AL, Nigam SK. Proteasome inhibition leads to a heat-shock response, induction of endoplasmic reticulum chaperones, and thermotolerance. J Biol Chem 272, 9086–9092 (1997).

48. Gamerdinger M, Kaya AM, Wolfrum U, Clement AM, Behl C. BAG3 mediates chaperone-based aggresome-targeting and selective autophagy of misfolded proteins. EMBO Rep 12, 149–156 (2011).

49. Rapino F, Jung M, Fulda S. BAG3 induction is required to mitigate proteotoxicity via selective autophagy following inhibition of constitutive protein degradation pathways. Oncogene 33, 1713–1724 (2014).

50. Wang H-Q, Liu H-M, Zhang H-Y, Guan Y, Du Z-X. Transcriptional upregulation of BAG3 upon proteasome inhibition. Biochemical and biophysical research communications 365, 381–385 (2008).

51. Ganassi M, et al. A Surveillance Function of the HSPB8-BAG3-HSP70 Chaperone Complex Ensures Stress Granule Integrity and Dynamism. Mol Cell 63, 796–810 (2016).

52. Du ZX, et al. Proteasome inhibitor MG132 induces BAG3 expression through activation of heat shock factor 1. J Cell Physiol 218, 631–637 (2009).

53. Lei Z, Brizzee C, Johnson GV. BAG3 facilitates the clearance of endogenous tau in primary neurons. Neurobiol Aging 36, 241–248 (2015).

54. Ji C, Tang M, Zeidler C, Hohfeld J, Johnson GV. BAG3 and SYNPO (synaptopodin) facilitate phospho-MAPT/Tau degradation via autophagy in neuronal processes. Autophagy 15, 1199–1213 (2019).

55. Cao YL, et al. A role of BAG3 in regulating SNCA/alpha-synuclein clearance via selective macroautophagy. Neurobiol Aging 60, 104–115 (2017).

56. Pegg AE. Functions of Polyamines in Mammals. J Biol Chem 291, 14904–14912 (2016).

57. Makletsova MG, Rikhireva GT, Kirichenko EY, Trinitatsky IY, Vakulenko MY, Ermakov AM. The Role of Polyamines in the Mechanisms of Cognitive Impairment. Neurochemical Journal 16, 283–294 (2022).

58. Freitag K, et al. Spermidine reduces neuroinflammation and soluble amyloid beta in an Alzheimer’s disease mouse model. J Neuroinffammation 19, 172 (2022).

59. Lewandowski NM, et al. Polyamine pathway contributes to the pathogenesis of Parkinson disease. Proc Natl Acad Sci U S A 107, 16970–16975 (2010).

60. Desiderio MA, Dansi P, Tacchini L, Bernelli-Zazzera A. Influence of polyamines on DNA binding of heat shock and activator protein 1 transcription factors induced by heat shock. FEBS Lett 455, 149–153 (1999).

61. Zhang S, et al. Acetylation of lysine 82 initiates TDP-43 nuclear loss of function by disrupting its nuclear import. bioRxiv, 2024.2009.2004.611121 (2024).

62. Neumann M, Rademakers R, Roeber S, Baker M, Kretzschmar HA, Mackenzie IR. A new subtype of frontotemporal lobar degeneration with FUS pathology. Brain 132, 2922–2931 (2009).

63. Nolan M, Talbot K, Ansorge O. Pathogenesis of FUS-associated ALS and FTD: insights from rodent models. Acta Neuropathol Commun 4, 99 (2016).

64. Urwin H, et al. FUS pathology defines the majority of tau- and TDP-43-negative frontotemporal lobar degeneration. Acta Neuropathol 120, 33–41 (2010).

65. Ormsby AR, Ramdzan YM, Mok YF, Jovanoski KD, Hatters DM. A platform to view huntingtin exon 1 aggregation flux in the cell reveals divergent influences from chaperones hsp40 and hsp70. J Biol Chem 288, 37192–37203 (2013).

66. Love MI, Huber W, Anders S. Moderated estimation of fold change and dispersion for RNA-seq data with DESeq2. Genome Biol 15, 550 (2014).

67. Brehme M, et al. A chaperome subnetwork safeguards proteostasis in aging and neurodegenerative disease. Cell Rep 9, 1135–1150 (2014).

68. Bordi M, De Cegli R, Testa B, Nixon RA, Ballabio A, Cecconi F. A gene toolbox for monitoring autophagy transcription. Cell Death Dis 12, 1044 (2021).

69. Stringer C, Pachitariu M. Cellpose3: one-click image restoration for improved cellular segmentation. Nat Methods 22, 592–599 (2025).

70. Zahavi EE, Ionescu A, Gluska S, Gradus T, Ben-Yaakov K, Perlson E. A compartmentalized microfluidic neuromuscular co-culture system reveals spatial aspects of GDNF functions. J Cell Sci 128, 1241–1252 (2015).

